# In vivo recellularization of xenogeneic vascular grafts decellularized with high hydrostatic pressure method in a porcine carotid arterial interpose model

**DOI:** 10.1101/2020.05.29.123018

**Authors:** Shunji Kurokawa, Yoshihide Hashimoto, Seiichi Funamoto, Akitatsu Yamashita, Kazuhiro Yamazaki, Tadashi Ikeda, Kenji Minatoya, Akio Kishida, Hidetoshi Masumoto

## Abstract

Autologous vascular grafts are widely used in revascularization surgeries for small caliber targets. However, the availability of autologous conduits might be limited due to prior surgeries or the quality of vessels. Xenogeneic decellularized vascular grafts from animals potentially substitute for autologous vascular grafts. Decellularization with high hydrostatic pressure (HHP) is reported to highly preserve extracellular matrix (ECM) which would be feasible for recellularization and vascular remodeling after implantation. In the present study, we conducted xenogeneic implantation of HHP-decellularized bovine vascular grafts from dorsalis pedis arteries to porcine carotid arteries then evaluated graft patency, ECM preservation and recellularization. Surgical procedure not to damage luminal surface of the grafts from drying significantly increased the graft patency at 4 weeks after implantation (P = 0.0079). After the technical improvement, all grafts (N = 5) were patent with mild stenosis due to intimal hyperplasia at 4 weeks after implantation. Neither aneurysmal change nor massive thrombosis was observed even without administration of anticoagulants nor anti-platelet agents. Elastica van Gieson and Sirius-red stainings revealed fair preservation of ECM proteins including elastin and collagen after implantation. Luminal surface of grafts was thoroughly covered with von Willebrand factor-positive endothelium. Scanning electron microscopy on luminal surface of implanted grafts exhibited cobblestone-like endothelial cell layer which is similar to native vascular endothelium. Recellularization of tunica media with alpha-smooth muscle actin-positive smooth muscle cells was partly observed. Thus, we confirmed that HHP-decellularized grafts are feasible for xenogeneic implantation accompanied by recellularization by recipient cells.

## INTRODUCTION

Cardiovascular disease is the highest cause of death worldwide. According to the report from world health organization, 17.9 million people died from cardiovascular diseases which accounts approximately 30 % of the whole death on 2016[1]. Coronary artery disease (CAD) and peripheral artery disease (PAD) are major components in cardiovascular disease with increased mortality and morbidity so far[2].

Revascularization surgery including bypass grafting is an established treatment modality for CAD and PAD. Considering relatively small size of target arteries, synthetic vascular prostheses made from expanded polytetrafluoroethylene, Dacron and others are not suitable for the revascularization of arteries with small caliber (< 4 mm) because of their poor patency rate even though their feasibility in large-(> 8 mm) or medium-(6 – 8 mm) caliber arteries[3–5]. Autologous grafts such as internal mammary arteries, radial arteries or saphenous veins are commonly used for bypass grafting in small caliber arteries instead. However, the availability of autologous arterial conduits is rather limited in general. Although autologous saphenous vein conduits possess higher availability and flexibility in regard of length and manipulation, it is reported that 20 – 45 % of patients requiring bypass surgeries to a small caliber targets have unavailable saphenous veins due to prior coronary or lower extremity revascularization, venous insufficiency, or prior vein surgeries[6, 7]. Alternative vascular conduits as a substitute for autologous grafts with appropriate tensile strength, viscoelasticity, biocompatibility, antithrombogenicity and biostability are anticipated.

Decellularization from xenogeneic tissue is reported to be a novel technology to prepare implantable tissues without immunologic problems caused by xenogeneic cellular components[8]. We have been investigating high hydrostatic pressure (HHP) method for decellularization[9] which can sufficiently deplete cellular components of the donor tissue with preserved extracellular matrices (ECM) proteins. The tissue tensile strength of decellularized grafts were superior to those using chemical detergents[10]. Xenogeneic decellularized vascular grafts with similar caliber of human autologous vascular grafts and with fair biocompatibility potentially address the unavailability of autologous grafts by virtue of the unlimited resources from animals.

In the present study, we investigated the feasibility and recellularization of HHP-decellularized vascular grafts as small-caliber vascular grafts in a xenogeneic porcine carotid arterial interpose model through histological, ultrastructural, morphological and hemodynamic evaluations of the implanted vascular grafts.

## MATERIALS AND METHODS

All experimental procedures were approved by the Kyoto University Animal Experimentation Committee (#17542) and performed in accordance with the guidelines for Animal Experiments of Kyoto University, which conforms to Japanese law and *the Guide for the Care and Use of Laboratory Animals* prepared by the Institute for Laboratory Animal Research, U.S.A. (revised 2011).

### Preparation of decellularized bovine dorsalis pedis artery

Edible bovine legs were obtained from a local slaughterhouse (Tokyo Shibaura Organ, Tokyo, Japan). The dorsalis pedis arteries were harvested from the legs and washed in saline. The arteries were packed in a plastic bag filled with saline and hydrostatically pressurized at 1000 MPa at 30 °C for 10 min using a cold isostatic pressurization machine (Dr. CHEF, Kobelco, Japan). The arteries were washed by continuous gradual shaking in saline supplemented with 0.2 mg/mL of DNase I and 50 mM of magnesium chloride for 7 days and then in 80 % ethanol for 3 days. After being washed, the arteries were stored in citric acid buffer (pH 7.4) at 4 °C until use.

### DNA quantification

The samples were freeze-dried and weighted (N = 5). The 20 mg of samples were minced and digested with 20 μg/mL proteinase K in 50mM of Tris-HCl, 1 % sodium dodecyl sulfate, 100 mM of NaCl, and 25 mM of ethylenediaminetetraacetic acid-2Na solution at 55 °C for overnight. DNA was isolated with a phenol/chloroform/isoamyl alcohol (25:24:1) extraction followed by ethanol precipitation. The amount of residual DNA was quantified by PicoGreen assay.

### Tensile test

The tensile test was conducted as previously described[10–12] using a universal testing machine (Autograph AG-X, Shimadzu) at crosshead speed of 0.1 mm/min. The samples were cut into dumbbell shape with a length of 35 mm and a width of 2 mm. The sample thickness was measured using micrometer with 2 μm accuracy before tensile test. Each specimen was preloaded to 0.01 N before loading. Four specimens from each group were separately tested. The elastic modulus was estimated from the slope of liner fit to the stress-strain curve. The tests were performed with longitudinal direction of the vessels.

### Animal model and surgical procedure of implantation

Nine to ten-month-old female CLAWN miniature swine (25 – 27 kg; N = 9) were anesthetized with ketamine (16 mg/kg body weight) and xylazine (1.6 mg/kg body weight) cocktail by intramuscular administration. Animals were intubated (Shily Hi-Lo oral tracheal tube 7.5mm I.D. COVIDIEN JAPAN, Tokyo, Japan) and maintained on 1 – 2 % isoflurane and 5 L oxygen using closed-circuit inhalation. During surgery, anesthesia depth and hemodynamic state were monitored by invasive blood pressure and electrocardiogram.

Decellularized grafts with HHP were implanted into right carotid artery. After midline neck incision, right common carotid artery was exposed for approximately 8 cm from carotid bifurcation. Heparin sodium (100 IU / kg) was administered before clamp of the artery. Right common carotid artery was interposed with HHP decellularized graft for approximately 4 cm in end-to-end fashion using 8-0 polypropylene continuous suture. In phase 1 group (N = 4), anastomoses were accomplished with similar surgical conditions with those in human bypass grafting surgery (conventional condition). In phase 2 group (N = 5), anastomosis was carefully performed keeping the decellularized graft, especially its luminal surface always wet with water (moist condition) and not touching its luminal surface. After anastomosis, blood flow was resumed. Blood flow velocity (time averaged maximum flow velocity) was measured by color Doppler ultrasound at the center of the implanted graft.

Wound was closed layer by layer. All animals were administered prophylactic cefazolin sodium intravenously. After implantation, no anticoagulants nor anti-platelet agents were administrated. Four weeks after transplantation, decellularized graft were explanted under general anesthesia and euthanized by intravenous bolus injection of potassium chloride (1 – 2 mEq/kg).

### Selective angiogram

Angiogram for bilateral carotid arteries are performed by inserting 6 Fr guiding catheter (INTRODUCER II, TERUMO, Tokyo, Japan) at femoral artery and selective angiogram with 4 Fr straight catheter (GLIDECATH, TERUMO) from the proximal end of common carotid arteries. Iopamidol (OYPLOMIN®300, FujiPharma, Toyama, Japan; 100ml) were used as contrast agent. The Angiogram were performed at just after implantation surgery, 2 weeks, 4 weeks after the surgery, respectively.

### Intravascular ultrasound (IVUS)

IVUS was performed at the same timing with that of angiogram, respectively. VISIONS PV .018 (Philips Japan, Tokyo, Japan) were used for IVUS transducer and VOLCANO (Philips Japan) were used for the data acquisition. IVUS was performed for bilateral carotid arteries (contralateral side of implanted side for control). The data were recorded from distal to proximal anastomosis site of the implanted graft to evaluate intraluminal stenosis and morphology. The data were analyzed by Image J software (version 1.50i, National Institutes of Health, Bethesda, MD)[13].

### Histological analysis

Explanted grafts were incised to longitudinal direction and fixed in 4 % paraformaldehyde for 48 hours. After fixation, explanted grafts were divided into proximal part and distal part, subsequentially embedded in paraffin. In each part, sections with 6 μm thickness were prepared consecutively and subjected to hematoxylin and eosin staining, Elastica van Gieson staining, Sirius red staining and von Kossa staining, respectively. Immunostaining of von Willebrand factor for endothelium (anti-von Willebrand Factor antibody, Abcam, Cambridge, UK, 1:3000) and α-smooth muscle actin for smooth muscle cells (anti-alpha smooth muscle actin antibody, Abcam, Cambridge, UK, 1:500) were performed, respectively. Sirius red staining sections were observed using polarized light microscope (BX51; Olympus, Tokyo, Japan).

### Scanning electron microscopy (SEM)

Native bovine dorsalis pedis arteries, decellularized grafts before implantation, native porcine carotid arteries and explanted grafts were fixed into a 2.5 % Glutaraldehyde 2 % Paraformaldehyde 0.1 M Phosphate Buffer solution (pH 7.4) for 24 hours at 4 °C, respectively. After fixation, samples were immersed into 1 % osmium tetroxide for 2 hours at 4 °C, and were dehydrated by graded ethanol (50 %, 60 %, 70 %, 80 %, 90 %, 95 % and 100 %) for 30 min. Subsequently, the samples were dried and coated with a thin layer of platinum palladium using an ion sputtering device (Eiko Corp.,Tokyo, Japan). The sample were examined with a Hitachi S-4700 scanning electron microscope (Hitachi, Tokyo, Japan).

### Statistical analysis

All data analyses were performed using JMP version 11.2.0 (SAS Institute, Cary, NC, USA). Statistical analysis of the data was performed with unpaired t-tests or Fisher's exact test for 2 groups. P < 0.05 was considered significant. Values are reported as means ± SD.

## RESULTS

### Physical and biochemical characteristics of decellularized vascular grafts with HHP method

Macroscopic view of decellularized bovine dorsalis pedis artery with HHP method is shown in Figure 1A. The amount of residual DNA was measured to quantify the efficiency of cell removal. The amount of residual DNA of decellularized graft was 15.85 ± 2.15 ng/mg which was significantly lower compared with those of bovine arteries, 4451.0 ± 351.7 ng/mg (P < 0.0001) (Fig. 1B).

**Figure 1:**
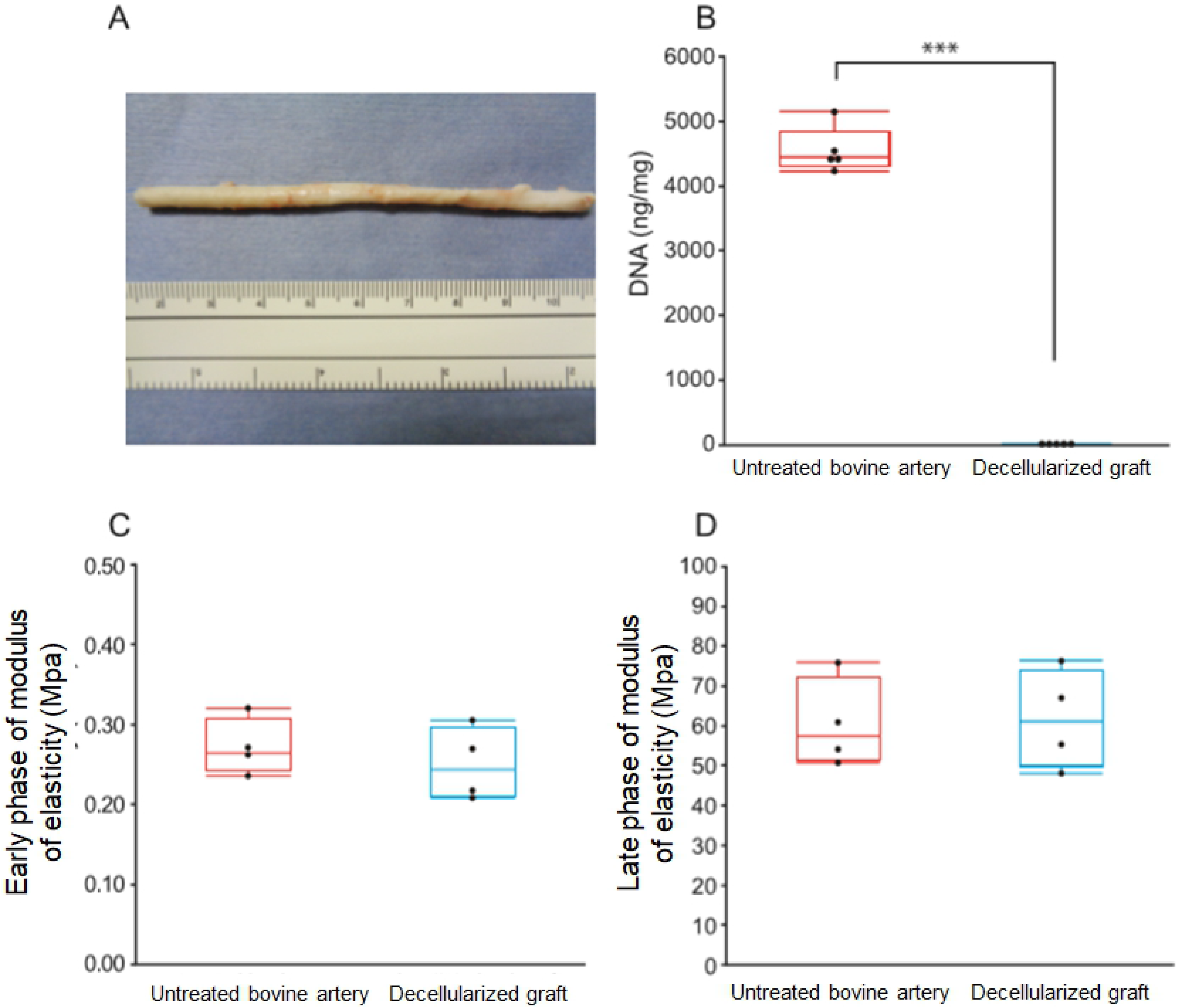
Characteristics of decellularized graft with HHP method. A representative macroscopic view of a decellularized bovine vessel with HHP method. HHP, high hydrostatic pressure. Upper scale of measure indicates centimeters, and lower indicates inchs, respectively. (**B**) Quantification of residual DNA. ***P<0.001. (**C**) Early phase of modulus of elasticity. (**D**) Late phase of modulus of elasticity.

Tensile test was performed to measure physical strength of decellularized grafts. Decellularized grafts exhibited 0.25 ± 0.05 MPa in early phase modulus of elasticity and 61.69 ± 12.49 MPa in late phase modulus of elasticity, while untreated bovine artery showed 0.27 ± 0.04 MPa and 60.42 ± 11.14 MPa (Fig. 1C, D), respectively. There are no significant differences in values of both phases (P = 0.50 in early phase modulus of elasticity, and P = 0.88 in late phase modulus of elasticity).

### Implantation surgery of HHP-decellularized vascular grafts

We started our experiments under surgically equivalent condition with that of usual human revascularization surgeries in preparation and anastomoses of autologous vascular grafts such as saphenous vein grafts (Phase 1; N = 4). We experienced that all 4 implanted grafts were occluded at 4 weeks after implantation (0/4; 0 % patency in Phase 1). We considered that the major reason of the occlusion might be the damage of the intima and exposure of basement membrane attributed by the drying of a lumen side which may affect antithrombogenicity and patency[14], then modified condition of anastomoses to moist condition not to allow lumen of grafts to be dried as much as possible (Phase 2; N = 5) (Fig. 2A,B; Supplemental video 1). In phase 2, we confirmed that all 5 grafts were patent at 4 weeks after implantation (5/5; 100 % patency in Phase 2) (Phase 1 vs Phase 2; P = 0.0079). There was no significant difference in time averaged maximum flow velocity just after anastomosis between Phase 1 and Phase 2 (Fig. 2C,D).

**Figure 2:**
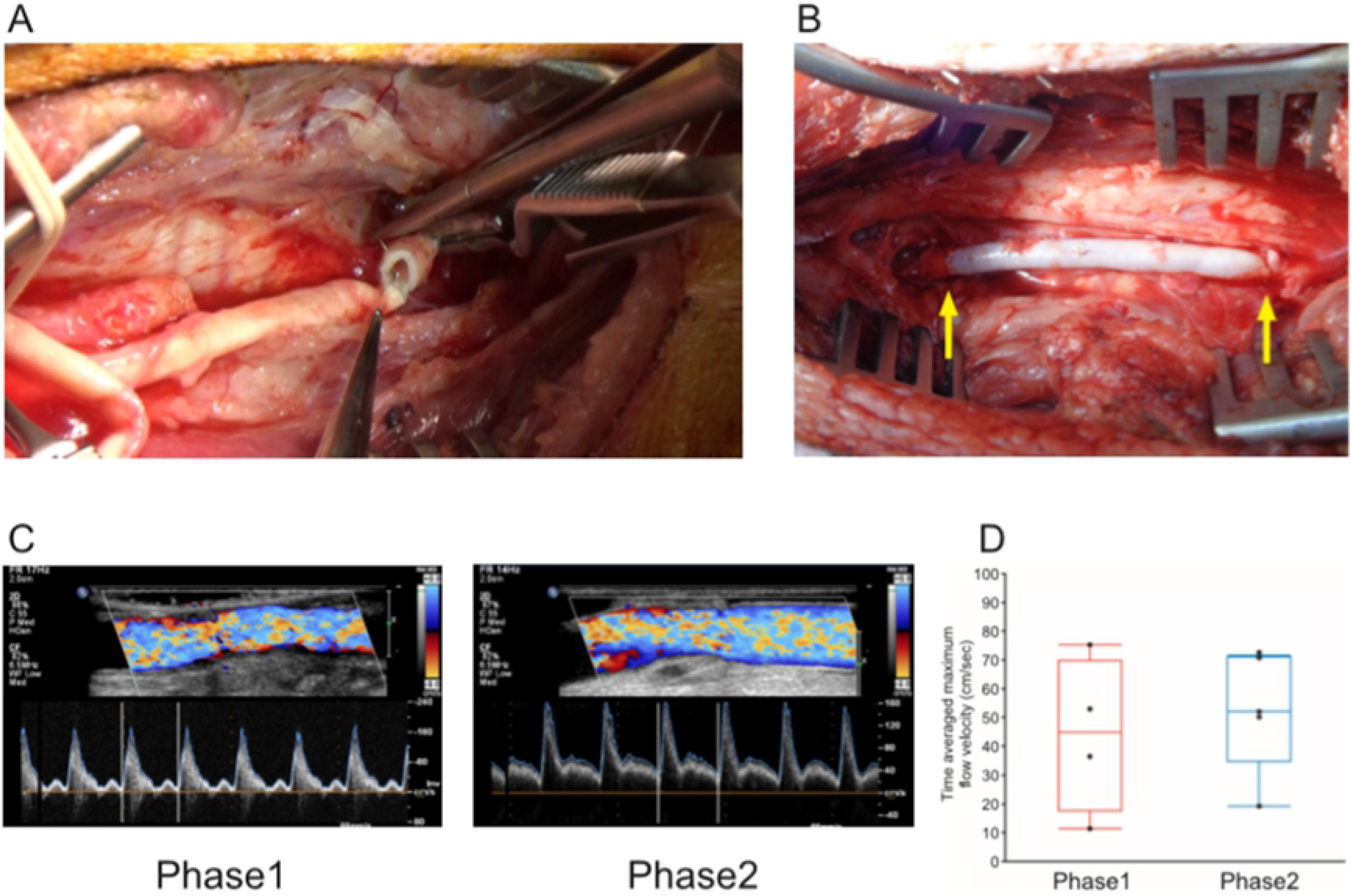
Implantation of decellularized graft into porcine carotid artery. (**A**) (**B**) A representative surgical view during (A) and after (B) anastomosis in moist condition (Phase 2). Yellow arrows indicate anastomosis sites. (**C**) Representative images of color Doppler ultrasound at the center of implanted graft (left; Phase 1, right; Phase 2). (**D**) Time averaged maximum flow velocity.

### Evaluations for implanted graft patency and morphology

In all cases of Phase 2, moderate stenoses of grafts were observed in the proximity of both anastomosis sites by selective angiogram (Fig. 3A). Intimal thickening of the corresponding regions was also confirmed by IVUS (Fig. 3B). Stenosis ratios of proximal and distal anastomotic region were 49.4 ± 0.12 % and 51.4 ± 0.04 %, respectively. We optimally evaluated all explanted grafts at 4 weeks after implantation. Luminal side of all explanted grafts were macroscopically smooth, but some explanted grafts were accompanied by small amount of thrombi (Fig. 3C). No aneurysmal change was observed in phase 2 cases.

**Figure 3:**
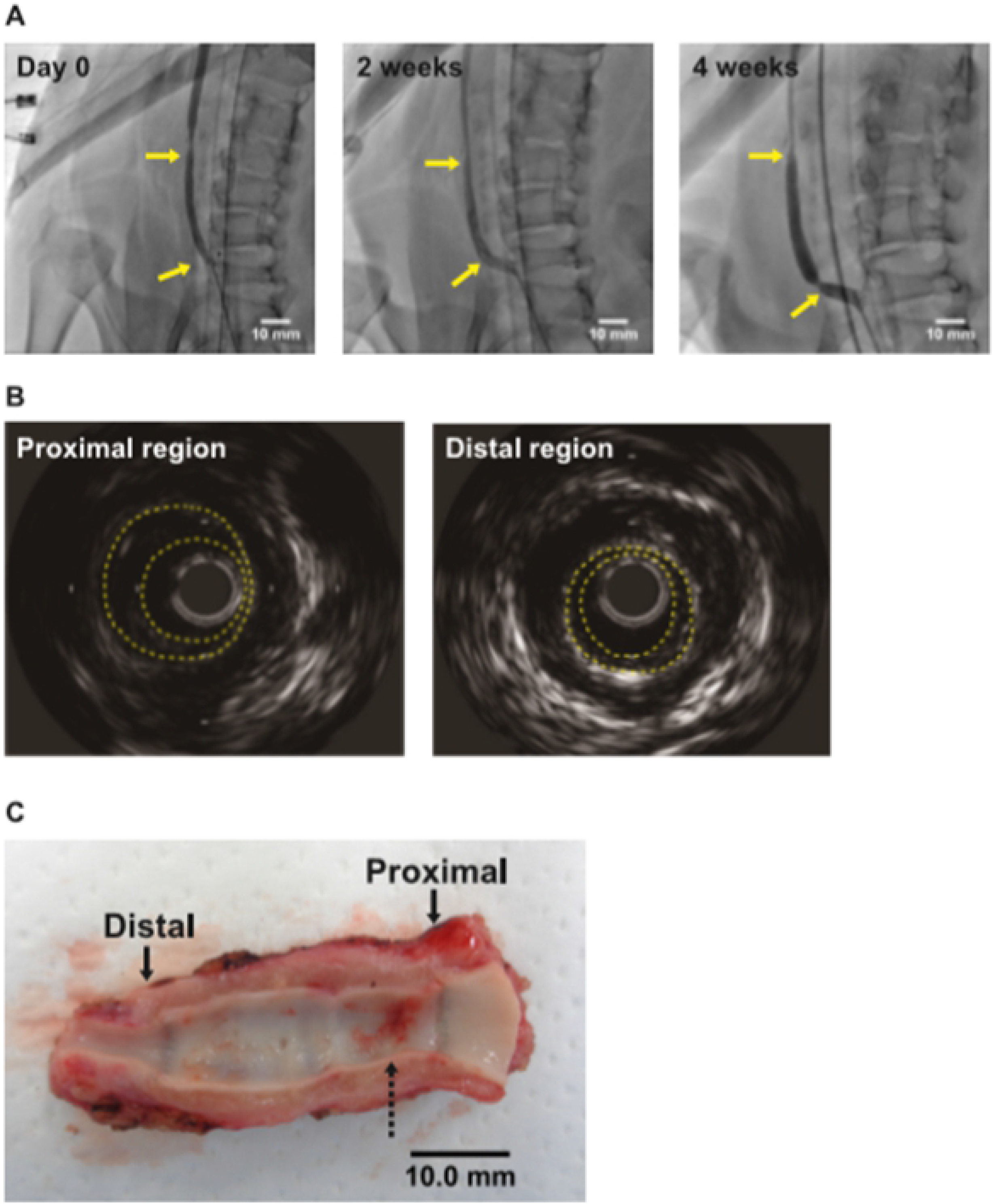
Evaluations for implanted graft patency and morphology. (**A**) Representative images of angiography of implanted grafts at just after implantation (left), and 2 weeks (middle) and 4 weeks (right) after implantation, respectively. Yellow arrows indicate anastomosis sites. (**B**) Representative images of IVUS. Left; proximal region, right; distal region. (**C**) A representative macroscopic view of explanted graft. Arrows indicate anastomosis sites. Dotted arrow indicates thrombi.

### Histological evaluations for implanted grafts after implantation

Hematoxylin and eosin staining revealed that HHP-decellularized graft did not contain cell nuclei indicating sufficient decellularization by HHP. On the other hand, recellularization in whole layers of the grafts was confirmed at 4 weeks after implantation (Fig. 4A). Elastica van Gieson staining and Sirius red staining exhibited a fair preservation of elastin layer (internal elastic lamina), tunica media consisted of collagen fibers and stratified elastin layers in HHP-decellularized grafts, and recellularization at 4 weeks after implantation, respectively (Fig. 4B). Polarized microscopical observations for striated elastin layer revealed that collagen I deposition was preserved among the elastin layers before implantation, and newly produced collagen III were deposited at the same region after implantation (Supplemental Fig. 1).

**Figure 4:**
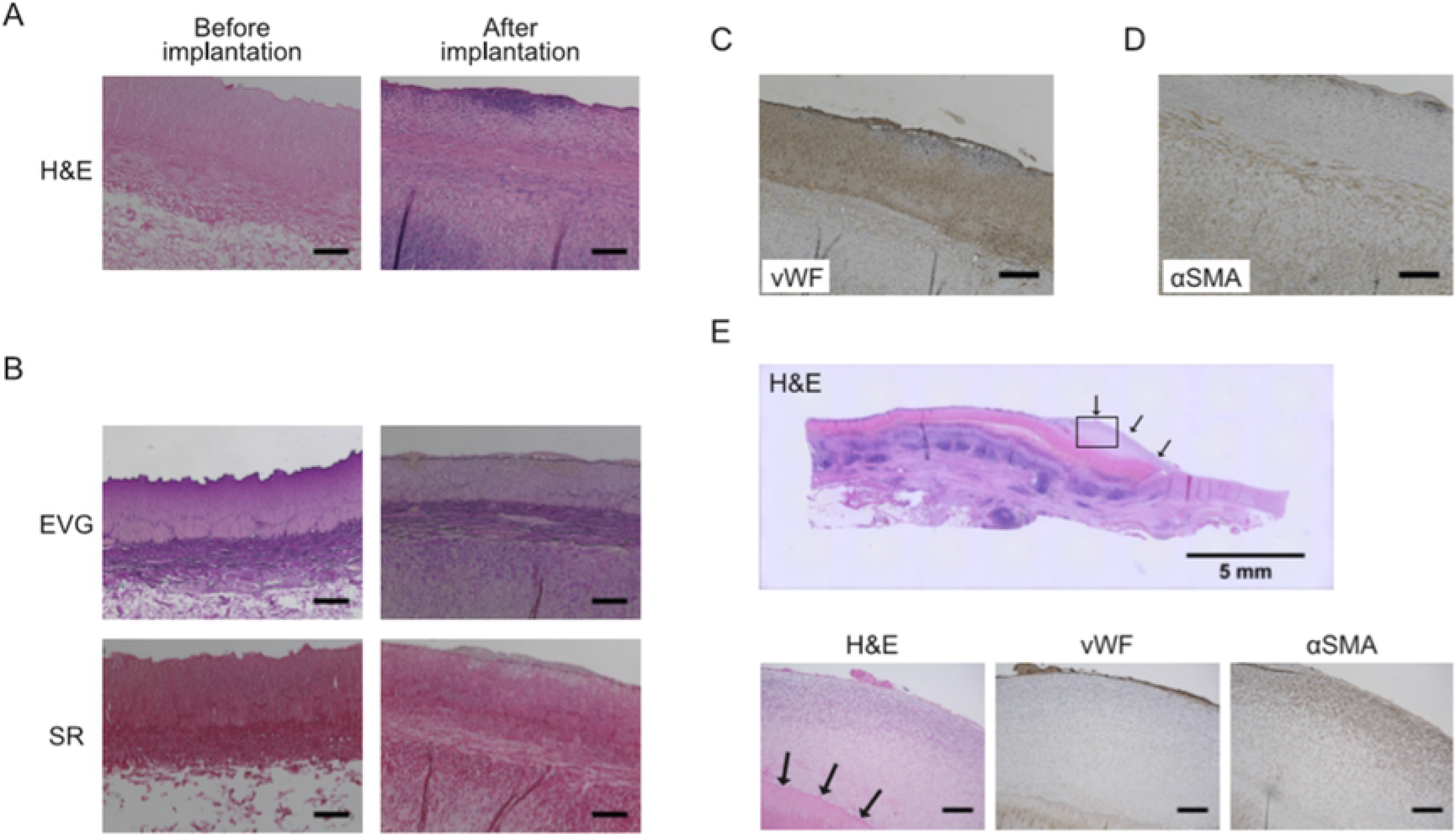
Histological evaluations for decellularized grafts after implantation (**A**) Representative Hematoxylin and Eosin (H&E) staining before implantation (left) and 4 weeks after implantation (right). Scale bars = 200 μm. (**B**) Representative Elastica van Gieson (EVG) staining and Sirius red (SR) staining, before implantation (left) and 4 weeks after implantation (right). Scale bars = 200 μm. (**C**) Representative von Willebrand Factor (vWF) immunostaining at 4 weeks after implantation. Scale bar = 200 μm. (**D**) Representative α-smooth muscle actin staining at 4 weeks after implantation. Scale bar = 200 μm. (**E**) H&E staining for representative stenotic regions close to proximal anastomoses at 4 weeks after implantation. Black arrows indicate hypertrophic region (top). H&E staining (bottom left), vWF immunostaining (bottom middle), and α-smooth muscle actin (αSMA) immunostaining (bottom right) for hypertrophic region indicated with square in top figure. Scale bars = 200 μm.

Immunostaining for von Willebrand Factor-positive endothelium revealed that the intima of implanted grafts was fairly covered by an endothelial cell layer throughout the graft (Fig. 4C). α-smooth muscle actin (αSMA)-positive vascular smooth muscle cells are observed among tunica media (Fig. 4D). These results indicate that the HHP-decellularized vascular grafts are recellularized by host-derived vascular cells in accordance with the anatomical allocations of native arteries.

We evaluated the stenotic regions in proximities of proximal and distal anastomoses. Hematoxylin and eosin staining exhibited that the hypertrophic regions are filled with proliferated cellular components. Immunostaining for von Willebrand Factor revealed thin endothelial cell layer covering the luminal surface. Immunostaining for αSMA showed that the stenotic region was mainly consisted of proliferated smooth muscle cells located between surface endothelial cell layer and internal elastic lamina (Fig 4E).

Von Kossa staining showed small deposition of calcified nodules close to the suture line which are not observed in another region of the graft (Supplemental Fig. 2). We evaluated the infiltration of inflammatory cells toward the implanted grafts. CD45 immunostaining revealed that the implanted grafts were not infiltrated with inflammatory cells except some regions close to luminal surface (Supplemental Fig. 3).

### SEM

The ultrastructure of luminal surface of vessels are evaluated by SEM. The luminal surface of native bovine artery (graft animal) was covered with endothelium exhibiting cobblestone-like appearance (Fig. 5A). After decellularization by HHP processes, the luminal surface of decellularized grafts exhibited acellular smooth surface without endothelium (Fig. 5B). The luminal surface of decellularized bovine graft implanted at porcine carotid artery for 4 weeks showed endothelium with cobblestone-like appearance by fair recellularization similar with those in bovine and porcine native arteries (recipient animal) (Fig. 5A, C, D).

**Figure 5:**
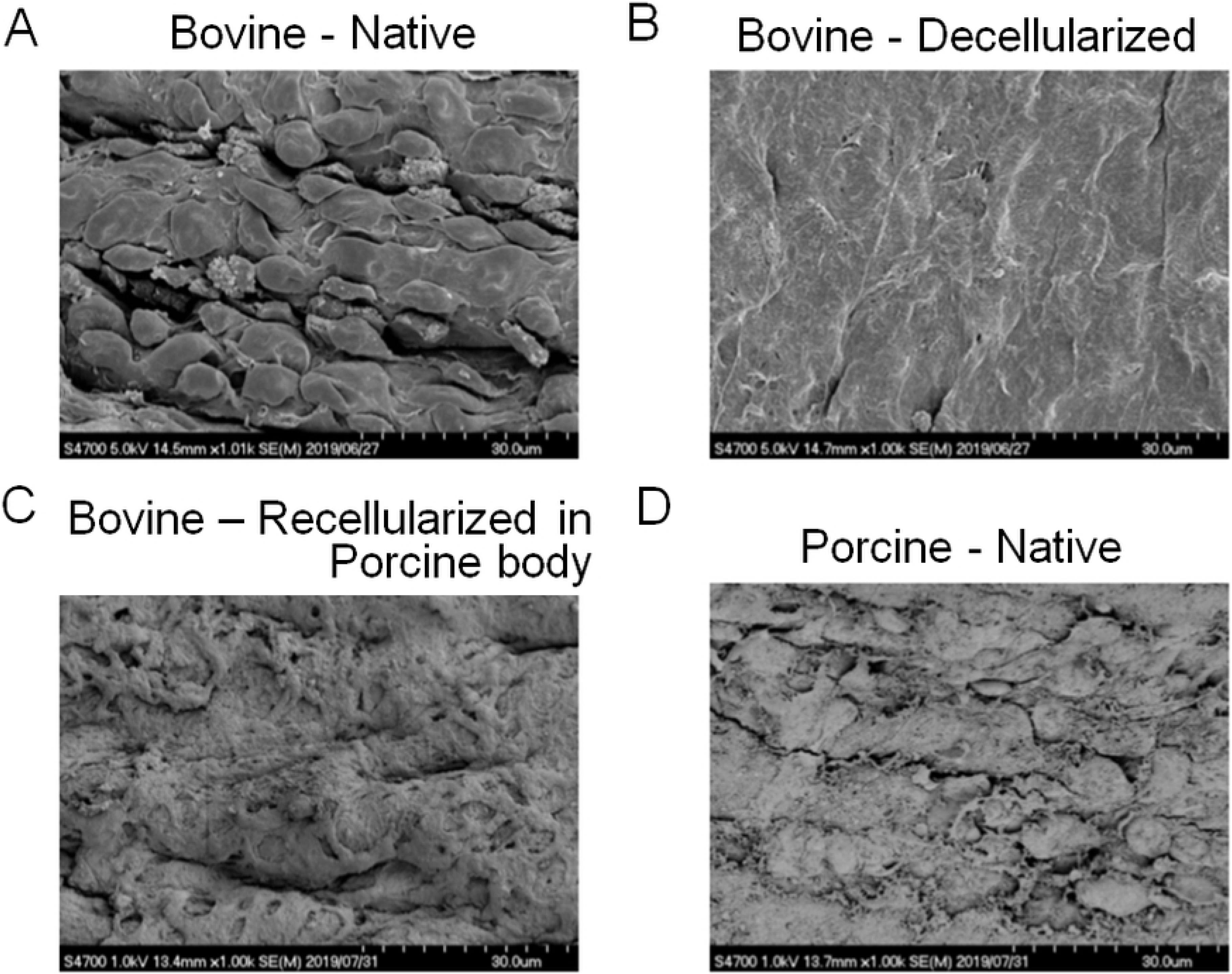
Evaluations for endothelial formation by scanning electron microscopy. (**A**) Representative luminal surface of untreated bovine artery. (**B**) Representative luminal surface of decellularized bovine artery. (**C**) Representative luminal surface of explanted graft. (**D**) Representative luminal surface of native porcine carotid artery

## DISCUSSION

In the present study, we confirmed that HHP-decellularized small-caliber vascular grafts can be recellularized by xenogeneic implantation with acceptable patency. Intimal layer of implanted grafts was covered by host-derived endothelial cells which could maintain antithrombogenicity of the graft. These results might indicate feasibility of the HHP-decellularized vascular grafts in xenogeneic implantation surgeries.

For patients requiring revascularization surgery, unavailability of autologous bypass grafts may lead to loss of an opportunity for appropriate therapy and consequent poor prognosis. Although cryopreserved arterial allografts can be a surrogate for autologous small-caliber grafts with acceptable long-term patency, the availability is rather limited because of the shortage of donors and insufficient operation of organ / tissue banks to preserve and provide allografts[15–17]. The present study was designed to address this healthcare problem through validating the feasibility of xenogeneic implantation of HHP-decellularized vascular grafts using bovine arterial grafts and a porcine carotid arterial interpose model.

Numerous methods for decellularization of living tissues have been reported so far[18–20]. In previous reports of the implantation of decellularized small-caliber vascular grafts by chemical or biological decellularization methods, insufficient outcomes such as graft occlusion, thrombus formation, and intimal proliferation throughout the graft were observed[21, 22]. We previously reported that the HHP decellularization, a novel method utilizing a physical basis can almost completely wash out the cellular components of porcine aorta and radial arteries with preserved mechanical properties such as elastic modulus[9]. In the present study, we decellularized bovine dorsalis pedis arteries by HHP and confirmed that the efficiency of cellular wash-out was >99% calculated by residual DNA amounts compared to those of untreated bovine dorsalis pedis artery. Residual DNA amounts in the present study (15.85 ± 2.15 ng/mg) satisfied the recommended criteria of successful decellularization as <50 ng/mg[23]. In mechanical tensile tests, there was no significant difference between HHP-decellularized and native arteries in the early and late elastic modulus representing the elastin phase and the collagen phase, respectively[24]. This result indicates that the physical strength of HHP-decellularized bovine arteries was not impaired by the HHP. Histological evaluations for HHP-decellularized arteries revealed that HHP did not damage the ECM structures of native arteries. Taken together, HHP method was proven to maintain structural and mechanical function of the arteries with enough depletion of cellular components.

The cellular source of *in vivo* recellularization in the present study would be recipient cells (not residual host cells) considering extremely high decellularization efficiency of the HHP method. Histological evaluations and SEM for decellularized grafts at 4 weeks after xenogeneic implantation revealed that the luminal surface was covered by the recipient's endothelial cells, and smooth muscle cells were infiltrated into the media layer. These results indicate that the recellularization took place as in an organized manner according to the original structure of the arteries. On the other hand, we simultaneously observed irregular structural reconstruction such as calcification around the suture line and intimal hyperplasia mainly consisted of smooth muscle cells proliferated at proximities of proximal and distal anastomoses. Even though the hyperplasia and calcification did not affect the blood flow of the implanted graft, histological changes and patency should be followed up for longer period in our future study.

In the present study, implantation surgeries were performed in 2 phases which were different in surgical conditions especially in moist conditions of the graft (Phase 1; conventional condition, Phase 2; no touch of luminal surface of the graft keeping moist condition). Flow patterns in 2 phases immediately after implantation did not differ each other, indicating that the qualities of anastomoses were not different in the 2 phases. However, the patency was significantly lower in Phase 1 compared to that in Phase 2 at 4 weeks after implantation. In Phase 1, all implanted grafts were resulted in thromboembolism, whereas all grafts in Phase 2 were patent with small amount of thrombi without any postoperative antiplatelet drugs or anticoagulants. These results suggest that the luminal surface of HHP-decellularized vascular grafts possesses fair antithrombogenicity when the intimal surface was not damaged by intraoperative grasping or drying. Although a careful manipulation would be required in bypass grafting surgeries, HHP-decellularized vascular grafts might hold promise as a novel vascular graft without antiplatelet agents or anticoagulants in the future.

## CONCLUSION

Xenogeneic HHP decellularized graft showed feasible capacity for recellularization and vascular remodeling without thrombogenicity. HHP decellularized vascular grafts may be utilized as new medical products for revascularization surgeries.

## ACKNOWLEDGEMENTS

This work was supported by research grants from the Ministry of Education, Culture, Sports, Science and Technology, Japan (to T.I.) (17H04291) and Supporting Program for Interaction-based Initiative Team Studies (SPIRITS), Kyoto University (to K.M.). The funders had no role in study design, data collection and analysis, decision to publish, or preparation of the manuscript. We thank Mr. S. Miyake, Ms. Y. Matsubara, Mr. H. Koda and Ms. K. Furuta (Kyoto University) for technical assistance.

The authors have declared that no competing interests exist.

